# Phenotype prediction in an *Escherichia coli* strain panel

**DOI:** 10.1101/141879

**Authors:** Marco Galardin, Alexandra Koumoutsi, Lucia Herrera-Dominguez, Juan Antonio Cordero Varela, Anja Telzerow, Omar Wagih, Morgane Wartel, Olivier Clermont, Erick Denamur, Athanasios Typas, Pedro Beltrao

## Abstract

Understanding how genetic variation contributes to phenotypic differences is a fundamental question in biology. Here, we set to predict fitness defects of an individual using mechanistic models of the impact of genetic variants combined with prior knowledge of gene function. We assembled a diverse panel of 696 *Escherichia coli* strains for which we obtained genomes and measured growth phenotypes in 214 conditions. We integrated variant effect predictors to derive gene-level probabilities of loss of function for every gene across strains. We combined these probabilities with information on conditional gene essentiality in the reference K-12 strain to predict the strains’ growth defects, providing significant predictions for up to 38% of tested conditions. The putative causal variants were validated in complementation assays highlighting commonly perturbed pathways in evolution for the emergence of growth phenotypes. Altogether, our work illustrates the power of integrating high-throughput gene function assays to predict the phenotypes of individuals.

**Highlights:**

- Assembled a reference panel of *E. coli* strains
- Genotyped and high-throughput phenotyped the *E. coli* reference strain panel
- Reliably predicted the impact of genetic variants in up to 38% of tested conditions
- Highlighted common genetic pathways for the emergence of deleterious phenotypes

## Introduction

Understanding the genetic and molecular basis of phenotypic differences among individuals is a long-standing problem in biology. Genetic variants responsible for observed phenotypes are commonly discovered through statistical approaches, collectively termed Genome-Wide Association Studies (GWAS, Bush and Moore 2012), which have dominated research in this field for the past decade. While such approaches are extremely powerful in elucidating trait heritability and disease associations (Yang et al. 2010; Welter et al. 2014), they often fall short in pinpointing causal variants, either for lack of functional annotation or due to lack of power to resolve variants in linkage disequilibrium (Edwards et al. 2013). Furthermore, by definition GWAS studies are unable to assess the impact of previously unseen or rare variants, which can often have a large effect on phenotype (Bodmer and Bonilla 2008). Therefore, the development of mechanistic models that address the impact of genetic variation on the phenotype can bypass the current bottlenecks of GWAS studies (Lehner 2013).

In principle, phenotypes could be inferred from the genome sequence of an individual by combining molecular variant effect predictions with prior knowledge on a gene’s contribution to the phenotype of interest. Such knowledge is now readily available by chemical genetics approaches, in which genome-wide knock-out (KO) libraries of different organisms are profiled across multiple growth conditions (Kamath et al. 2003; Dietzl et al. 2007; Hillenmeyer et al. 2008; Nichols et al. 2011; Price et al. 2016). One of the outputs of such screens are genes whose function is essential for growth in a given condition. Variants negatively affecting the function of those genes should likely be associated with individuals displaying a significant growth defect in that same condition. As the impact of variants on gene function can be inferred using different approaches (Thusberg, Olatubosun, and Vihinen 2011; Kulshreshtha et al. 2016), such mechanistic models offer straightforward molecular explanations of their impact. As rare or previously unseen variants can also be used by this approach, there lies the possibility to deliver predictions of phenotypes at the level of the individual. Combining variant effector predictors with gene KO information was previously tested with some success in the budding yeast *Saccharomyces cerevisiae*, but only on a limited number of individuals and conditions (15 and 20, respectively, Jelier et al. 2011). Given the growing availability and investment into genome-wide gene functional studies, there is an opportunity to apply such an approach more extensively.

In contrast to eukaryotes, diversity within the same bacterial species can result in two individuals differing by as much as half of their genomic content. *Escherichia coli* is not only one of the most studied organisms to date (Blount 2015), but also one of the most genetically diverse bacterial species (Lukjancenko, Wassenaar, and Ussery 2010; Tenaillon et al. 2010). Individuals of this species (termed "strains"), exhibit a diverse range of genetic diversity, from highly homologous regions to large differences in gene content, collectively termed "pan-genome" (Medini et al. 2005). Phenotypic variability is therefore likely to arise from a combination of single nucleotide variants (SNVs) and changes in gene content. Since conditional essentiality has been heavily profiled for the reference *E. coli* strain (K-12) (Nichols et al. 2011; Price et al. 2016, Herrera-Dominguez et al., unpublished), we set out to systematically test the applicability of such genotype-to-phenotype predictive models for this species. We reasoned that it would also test the limits of the underlying assumption of such models, which is that the effect of the loss of function of a gene is independent of the genetic background (Dowell et al. 2010).

We therefore collected a large and diverse panel of E. coli strains (894), for which we measured growth across 214 conditions, as well as obtained the genomic sequences for the majority of the strains (696). For each gene in each sequenced strain we calculated a "gene disruption score” by evaluating the impact of non-synonymous variants through conservation and structural analysis. We then applied a model that combines the gene disruption scores with the prior knowledge on conditional gene essentiality to predict phenotypes across strains in a condition specific manner. The model yielded significant predictive power for 38% of conditions having at least 5% of strains with growth defects. We independently validated a small number of causal variants with complementation assays. Since our predictions did not apply equally well for all conditions, we conclude that the set of conditionally essential genes has diverged substantially across strains. Overall, we anticipate that this *E. coli* reference panel presented here will become a community resource to address the multiple facets of the genotype to phenotype research, ultimately enabling the development of biotechnological and personalized medical applications.

## Results

### An *E. coli* strain collection for genotype-to-phenotype analysis

We have assembled a large genetic reference panel of *E. coli* strains, able to capture the genetic and phenotypic diversity of the species. The collection comprises 894 *E. coli* strains, broadly divided into natural isolates (527 strains) and strains derived from experimental evolution experiments (367 strains). The natural isolates can be further divided into laboratory strains (16 strains), commensals (330 strains), pathogens (153 strains) or others (28 strains). The collection contains widely studied laboratory and pathogenic isolates, including BW25113 (Datsenko and Wanner 2000) (background of the KEIO knockout collection, Baba et al. 2006), MG1655 and W3110 (classical K-12 model strains, Bachmann 1972), DH5a (broadly used for cloning) and many model pathogenic strains (UPEC, EPEC, EAEC and APEC). The strain collection also includes the ECOR (72 strains, Ochman and Selander 1984), IAI (82 strains, Picard et al. 1999) and NILS (82 strains, Bleibtreu et al. 2014) strain collections, as well as 290 strains from the LTEE collection (E. *coli* Long Term Evolution Experiment) (Tenaillon et al. 2016). The collection additionally includes 15 strains belonging to other species of the *Escherichia* genus. The full list of strains, including name, collection of origin and links to relevant databases is provided in Supplementary table 1 and online at https://evocellnet.github.io/ecoref. Such a large and diverse collection of strains serve as the foundation of the development of phenotypic predictive models.

Our collection also exhibits a high diversity at the genetic level. We combined already available genomic sequences (374) with newly generated ones (322) to obtain the genomes of 696 strains, after removing strains whose sequence did not match known typing information and duplicate isolates (Figure 1A and Supplementary table 1 and Methods). When compared to the reference strain *E. coli* K-12 we observed an expected linear relationship between core genome phylogenetic distance and number of nonsynonymous mutations (Supplementary figure 1); we also observed a similar linear relationship with the number of reference genes missing in each strain (Supplementary figure 1), with up to 587 reference genes missing in strain NR-15878. We found a similar behaviour for the number of additional genes present in each strain compared to the reference, with strain DE-C0MM-4000 having 2178 genes not present in K-12. Such genetic plasticity is a known feature of bacterial pan-genomes (Medini et al. 2005), and the so-called accessory genome (i.e. genes present only in a subset of individuals) can significantly factor in determining phenotypes (Brynildsrud et al. 2016).

**Figure 1.**
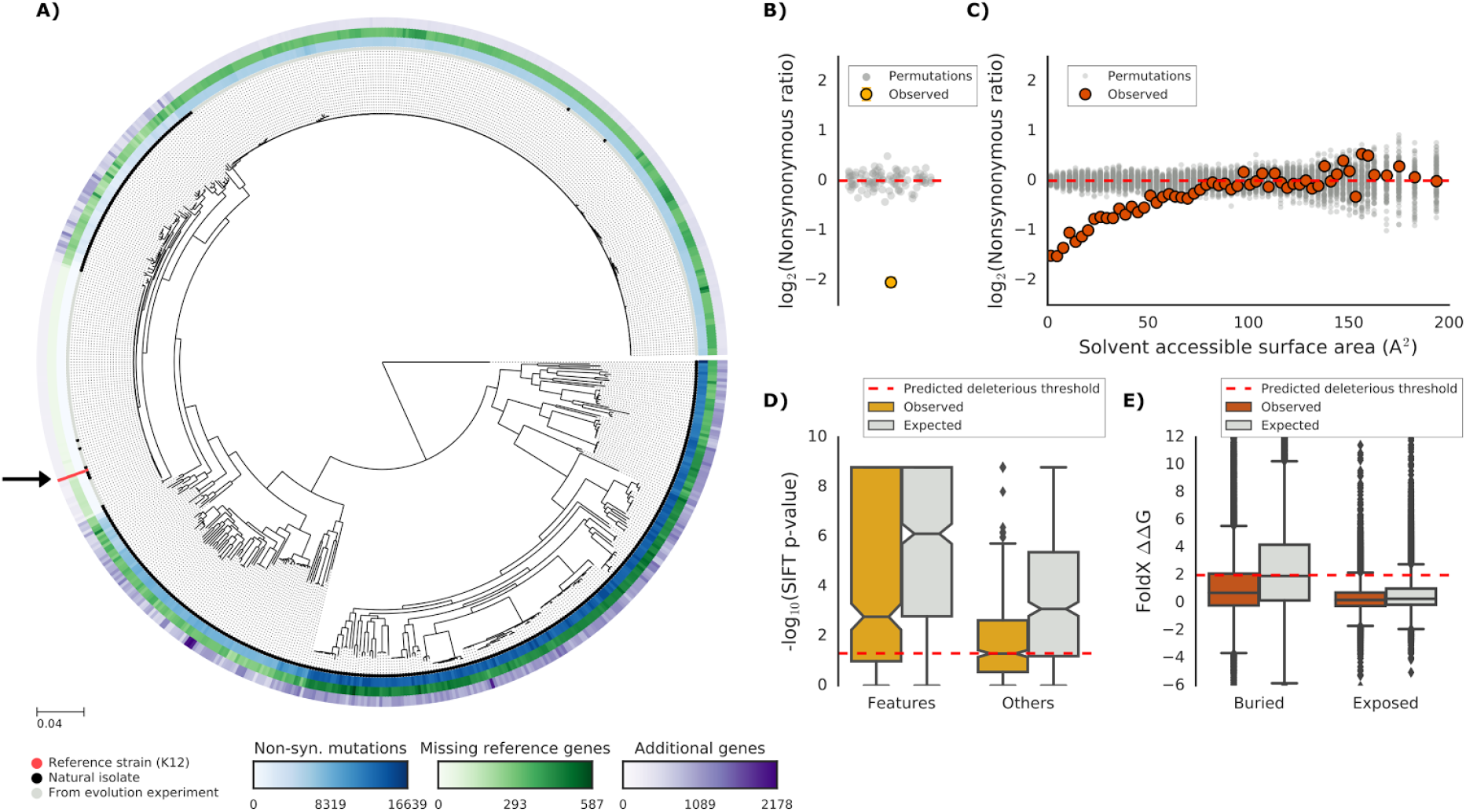
Genetic variability and protein sequence constraints in the *E. coli* strain collection. **(A)** Core genome SNP tree of the members of the strain collection; shades of blue in the inner ring indicate the number of nonsynonymous substitutions with respect to the reference strain (E. *coli* K-12), shades of green in the middle ring indicate the number of reference genes absent from each strain and shades of purple in the outer ring indicate the number of additional genes present in each strain when compared to reference strain. Black arrow indicates the reference strain. **(B-C)** The nonsynonymous substitutions ratio (proportion of observed mutations in selected site over a random sample of all other sites) across the collection proteomes is used to highlight the presence of evolutionary constraints, through a comparison against shuffled sites positions. **(B)** Functional sites in proteins (derived from UniProt(UniProt Consortium 2015) annotations) show a lower density of nonsynonymous substitutions than the rest of the protein. **(C)** Residues with lower predicted water accessibility show a lower density of nonsynonymous substitutions than residues with higher accessibility to the solvent. **(D-E)** Comparison of the predicted impact of observed nonsynonymous substitutions in constrained sites against all other sites, and comparison with the impact of all possible nonsynonymous substitutions (a random set of the same size). **(D)** Predicted impact of nonsynonymous substitutions, as measured by the SIFT (Ng and Henikoff 2001) algorithm, shows that substitutions in functional constrained sites are more benign than expected. **(E)** Predicted impact of nonsynonymous substitutions, as measured by the FoldX (Guerois, Nielsen, and Serrano 2002) algorithm, shows that substitutions in structurally constrained sites are more benign than expected.

### Functional constraints in the *E. coli* proteome

The first step in predicting the impact of genetic variability is to assess its molecular and cellular consequences. It is expected that the majority of variants will be neutral, as deleterious mutations should be counter-selected. Moreover, nonsynonymous substitutions can be assumed to be the strongest contribution to phenotypic variability, particularly in bacteria, where most of the genome encodes for proteins (Lynch and Conery 2003). Such variants can affect a protein either by altering its function or its structural stability, leading to a loss of function or cytotoxic effects. We therefore tested whether functionally and structurally important residues are under purifying selection in this strain collection by measuring the frequency of nonsynonymous variants at these important regions compared to a random background. Clear signs of purifying selection were evident for sites that are important for protein function and structure. For both functional sites (e.g. enzymes active sites, metal binding sites) and residues buried inside the protein’s structure we observed a lower density of nonsynonymous substitutions when compared to the other sites (Figure 1B and 1C).

We then asked if variant effect predictors are capable of inferring the importance of these functional regions. For this we derived structural models and protein alignments covering 60.2% and 94.7% of the *E. coli* K-12 proteome, respectively (or 50.9% and 95.9% of all protein residues) and used them to compute the impact of all possible nonsynonymous variants (see Methods, available at http://mutfunc.com, Wagih et al., unpublished). Reflecting the clear evolutionary constraints acting on the *E. coli* proteome, the predicted impact of naturally occurring substitutions in functionally and structurally buried residues is significantly lower than for random mutations in the same sites (Figure 1D and 1E). Thus, not only important regions in the protein are more conserved, but also the impact of substitutions occurring in these regions is more benign than the predicted impact of random variants.

### Gene level predictions of variant effects

The impact of all individual variants within a gene can be combined to predict whether its function is likely to be affected. We leveraged the approach developed in S. *cerevisiae* (Jelier et al. 2011), where the predicted impact of each nonsynonymous substitution is combined into a single likelihood measure of gene disruption (termed here "gene disruption score", Figure 2A and Methods), using a multiplicative model. We also included a simple heuristic to account for reference genes missing from specific strains, by assuming that their function is completely impaired (maximum gene disruption). The absence of a gene does not exclude the presence of another gene that could compensate for the lost function. However, predicting the function of members of the accessory genome is a notoriously difficult problem (Radivojac et al. 2013), and we therefore did not consider that part of the pan-genome.

**Figure 2.**
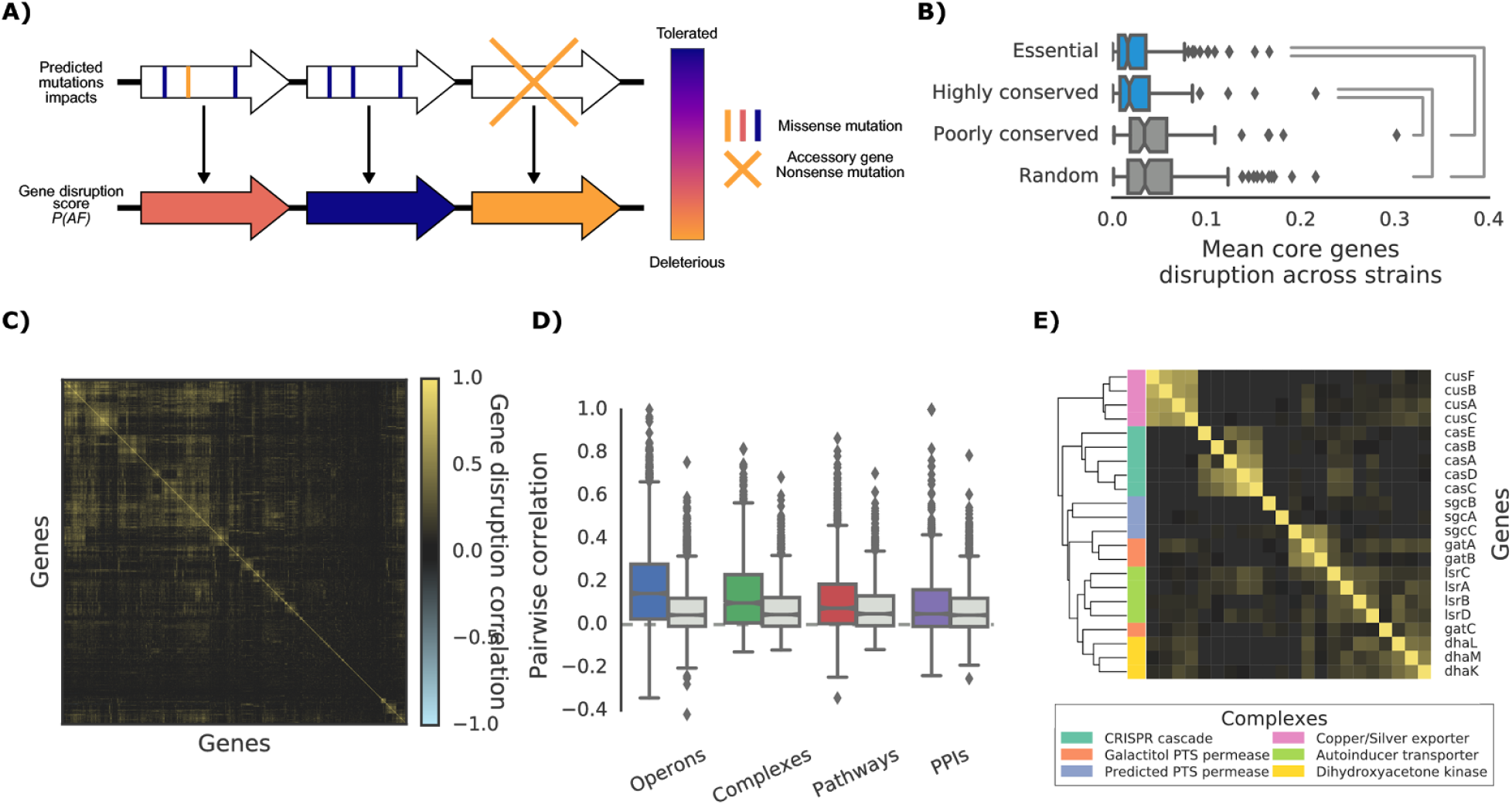
The “gene disruption score”, a gene-level prediction of the impact of nonsynonymous substitutions, and its biological properties. **(A)** Schematic representation of how all substitutions affecting a particular gene are combined to compute the gene disruption score. With the term “accessory gene” we indicate those genes that are present in the reference strain but not in the focal one. **(B)** Average gene disruption score across all strains for four categories of genes conserved in all strains (“core genome”). Essential genes are retrieved from the OGEE database (Chen et al. 2012), highly (or poorly) conserved genes are defined as those genes found in more (or less) than 95% of the bacterial species present in the eggNOG orthology database (Huerta-Cepas et al. 2016). Statistically significant differences (Cohen’s d value > 0.3) are reported. **(C)** Gene-gene correlation profile of gene disruption across all strains shows clusters of potentially functionally related proteins. **(D-E)** The gene disruption correlation profiles as a predictor of genes functional associations. **(D)** Gene-gene pairwise gene disruption correlation inside each annotation set (colored boxes) and inside a random set of genes of the same size (grey boxes). **(E)** Gene-gene correlation profile of gene disruption in protein complexes with high disruption score correlation.

We then examined if the calculated gene disruption score can be considered as a relevant measure of the impact of mutations on gene function by correlating it with genetic indicators of high functional importance. Essential and phylogenetically conserved genes show a lower average gene disruption score across all strains, as compared to less conserved or random ones (Figure 2B). This is expected, as both categories are more likely to have stronger evolutionary constraints and should, therefore, maintain their function inside the species.

We also probed whether genes that are predicted to lose their function together across all strains are functionally associated. To test this, we used the gene disruption score correlation profile across all strains (Figure 2C) as a predictor for known gene functional associations, such as operons, protein complexes, functional pathways and protein-protein interactions (PPIs). Pairwise gene disruption correlation is indeed significantly higher between functionally associated proteins than between random pairs of genes (Figure 2D), and therefore it can be used as a predictor of functional modules (Supplementary figure 2).

Some examples of protein complex subunits with highly correlated gene disruption scores are provided in Figure 2E. Those include the CRISPR cascade complex (*casABCDE*), a copper/silver transporter (*cusABCF*), an autoinducer transporter (*IsrABCD*), a dihydroxyacetone kinase (*dhaLKM*), and two phosphotransferase (PTS) permeases (*gatABC* and *sgcABC*). In some cases, we observed a strong correlation among all members of the protein complex, such as *cusABCF*, while in others the correlation is limited to only a few members, such as *gatABC*, where only *gatA* and *gatB* are predicted to be co-affected. The *sgcC* transmembrane subunit of the *sgcABC* predicted PTS permease appears to be highly correlated with *gatA* and *gatB*. As *sgcC* is a known homolog to *gatC*, we could speculate on a putative functional interaction between *gatAB* and *sgcC*. Some of the strongest correlations are driven almost exclusively by nonsynonymous variants that are predicted to be deleterious, such as in *cusABCF*, which is present in ~95% of the strains. In other cases, the correlations are driven both by the complex being co-lost or co-mutated, such as *casABCD*. When partitioning the reference genes into “core” (conserved in all strains) and “accessory” (present only in a subset of strains), co-mutation patterns are more predictive of functional associations in accessory genes, even when computing the gene disruption score from nucleotide substitutions only (Supplementary figure 2). This is presumably due to stronger evolutionary constraints on core genes, leading to a reduction in the correlation signal due to few instances of genes losing their function significantly. Altogether, our results point to functional modules being co-mutated or co-lost across strains of the same species, suggesting a very fast local adaption to specific conditions, similar to that observed in pathogens restricting their niche/host specificity (Reuter et al. 2014).

Taken together, these observations suggest that the gene disruption score is a biologically relevant measure of the impact of mutations at the gene level and that it could be used for growth phenotype predictions.

### The phenotypic landscape of the *E. coli* collection

To explore the applicability and performance of genotype-to-phenotype predictive models we tested the whole strain collection (909 strains) fitness on a large (214) variety of conditions. The conditions (Supplementary Table 2 and 3) included chemical substances (such as antibiotics), environmental stressors (such as high temperature or exposure to UV light), and a variety of nutrient sources and conditioned media (e.g. supernatants of other bacteria). To profile all these conditions, we used high-density colony arrays and measured colony size as a proxy for fitness, with the same experimental setup used before for the *E. coli* K-12 knockout (KO) library (Nichols et al. 2011). The majority of the conditions (161 out of 214) were tested at the same time and with the same laboratory conditions as the K-12 KO library (Herrera-Dominguez et al., unpublished), thus reducing experimental variability between the fitness measurements for the strains and the K-12 KO collection.

We used the deviation of each strain’s colony size from itself across all conditions and all other strains in the same condition as our final phenotypic measure (termed “S-score”, Collins et al. 2006). Thereby we obtained a list of conditions for which we know whether a tested strain has grown significantly more or less than the expectation (Figure 3A and Methods). Both biological replicates (Figure 3C) and strains present in two distinct plates (Supplementary figure 3) were highly correlated (Pearson’s r 0.693 and 0.648, respectively), indicating that we measured phenotypes with high confidence. The full phenotypic matrix for each strain across all conditions contains 114,004 single measurements (Supplementary material 1).

**Figure 3.**
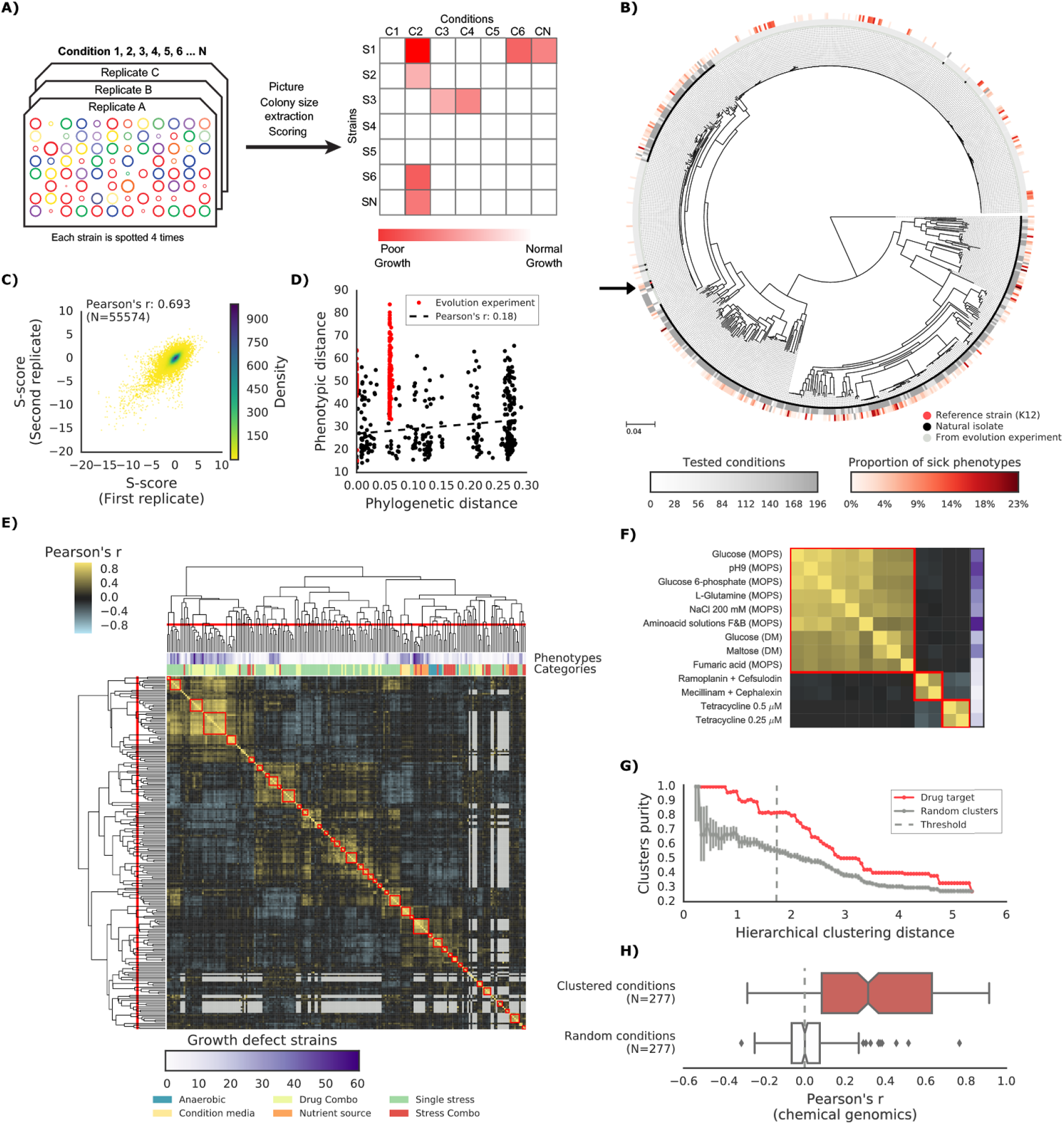
The phenotypic landscape of the *E. coli* strain collection. **(A)** Phenotypic screening experimental design and data analysis. Three plate replicates (biological) are screened per condition, with each strain being present in at least four copies on each plate. Colony sizes across all conditions are then used to compute an S-score (Collins et al. 2006) for each strain in each condition, capturing the strain fitness effect in this condition. **(B)** Core genome SNP tree for all strains in our collection. Grey shades in the inner ring indicate the number of conditions tested for each strain, red shades in the outer ring indicate the proportion of tested conditions in which the strain shows a growth phenotype (strain grows significantly less than expected). Black arrow indicates the reference strain. **(C)** Phenotypic measurements show good replicability, as measured by pairiwise comparing the S-scores of the three biological replicates and between strains present in multiple plates. **(D)** Person’s correlation between phylogenetic and phenotypic distance. **(E)** Hierarchical clustering of condition correlation profiles; the threshold is defined as the furthest distance at which the minimum average Pearson’s correlation inside each cluster is above 0.3. The two colored bands on top indicate the number of strains with growth defects for each condition and its category, showing consistent clustering. Gray-colored cells in the matrix represent missing values due to poor overlap of strains tested in the two conditions. **(F)** Detailed view of three clusters highlighted in panel e. **(G)** Clusters purity (computed for drug targets) for each hierarchical distance threshold, against that of random clusters (100 repetitions) shows that drugs with similar target tend to cluster together. **(H)** Condition pairwise correlation in the *E. coli* K-12 chemical genomics data is significantly higher for the condition clusters defined by the phenotypic screening than for a random set of condition clusters of equal size.

As opposed to genetic variability, we detected no strong positive correlation between phylogenetic and phenotypic distance – both calculated relative to reference K-12 strain (r=-0.33, p-value=3E-19, Figure 3D). A very weak positive correlation was evident when excluding the highly similar strains from evolutionary experiments (r=0.18, p-value=0.0007, Figure 3d). We also did not observe a strong correlation between the phylogenetic distance to K-12 and the fraction of growth phenotypes (Supplementary figure 3). These findings reinforce the idea that most variants across these stains are neutral. For instance, strain ECOR-12, despite being relatively similar in DNA sequence to the K-12 reference strain (1,942 nonsynonymous substitutions and 191 reference genes missing), grows poorly in 21 conditions, whereas strain ECOR-53, one of the strains with the highest distance to K-12 (13,490 nonsynonymous substitutions and 449 reference genes missing), is able to grow in all tested conditions.

Of particular interest are those strains derived from evolution experiments, such as the members of the LTEE collection (Tenaillon et al. 2016). While most strains grow in all tested conditions (189 strains out of 266 tested), a significant fraction (77) showed at least one growth phenotype. Again, the phylogenetic distance from the parental strains (REL606 and REL607) is not correlated with the proportion of growth phenotypes (Pearson’s r: 0.08), even though hypermutators exhibit a slightly higher number of phenotypes (Cohen’s d: 0.513). These results clearly underline the large phenotypic space covered by the *E. coli* strain collection and that simple metrics of phylogenetic similarity are not predictive of phenotype differences. Instead, few DNA variants are sufficient to cause clear phenotypic differences, indicating the importance of statistical or predictive strategies to prioritize those variants.

We observed a significantly higher sensitivity (Cohen’s d: 0.651) of pathogenic strains (N=149) when compared to commensal ones (N=344), as measured by the proportion of tested conditions in which each strain shows a growth phenotype (Supplementary figure 3). Such observation is consistent with previously observed lower metabolic capabilities for virulent strains of *E. coli* (Durso, Smith, and Hutkins 2004).

To test the accuracy of the fitness measurements, we used the correlation of phenotypic profiles across all strains to derive groups of similarly behaving conditions across the strains (Figure 3E). Condition’s clustering was consistent with the conditions macro-categories (e.g. stresses *versus* nutrient sources) and with the number of sensitive strains. For example, most of the conditions related to alternative nutrient sources elicited similar fitness outcomes across strains. The correlation of phenotypic profiles also clustered drugs with same mode of action (MoA), as shown by comparing the cluster’s purity against those of random clusters (Figure 3G).

As an additional benchmark for the accuracy of the fitness responses we tested if pairs of drugs that have correlated fitness effects across our strain collection also displayed correlated profiles in the K-12 knock-out collection of strains. Indeed, those conditions that clustered together in our data also had higher correlations in the *E. coli* K-12 knockout library than random pairs of conditions (Figure 3H). Using principal component analysis we concluded that the phenotypic space sampled by the strain collection is largely similar to that of the K-12 KO library (Supplementary figure 3). This suggests that the phenotypes exhibited by the natural strains are similar to those reached by the K-12 gene KO, even if the strains possess much larger genetic variability in gene content and nonsynonymous variation.

We have generated a rich phenotyping resource for our *E. coli* reference strain panel. This resource is credible as it recapitulates known biology, and it surveys a rich phenotypic space, providing insights into the evolution of phenotypes within the species.

### Predictive models of conditional growth defects in the *E. coli* strain collection

To build phenotype prediction models for all strains in the strain collection, we combined the gene disruption scores with information on the K-12 conditionally essential genes. We then computed a conditional score that would rank the strains according to their predicted growth level in the tested condition, from normal growth to the most defective growth phenotype. This was done for the 148 conditions which had at least one strain displaying growth defects in our screen and had also been tested in the *E. coli* K-12 KO collection. This ranking was then compared to the determined phenotypes for all *E. coli* strains and the Area Under the Curve of a Precision-Recall curve (PR-AUC) was used for assessing our predictive power (Figure 4A and Methods). Since gene essentiality can be influenced by the genetic background, and therefore change between strains (Dowell et al. 2010), we weighted the contribution of each conditionally essential gene according to their level of essentiality and functional importance reasoning that highly important genes would more likely have conserved functions across stains (Methods). Our predictive score is able to discriminate strains with normal growth from ones with growth defects with significantly higher power than randomized scores. Both the predicted impact of single nucleotide variants and gene presence/absence patterns contribute to the predictive power of the model (Supplementary figure 4). The predictive power increases for conditions in which more strains displayed growth defects (Figure 4B). We found a significant correlation between predicted and measured growth defects for 20% of conditions that have at least 1% of poorly growing strains. The predictive capacity is higher for conditions with larger number of strains with growth defects (Supplementary figure 4), reaching 38% for all conditions with more than 5% of poorly growing strains. No class of condition (antibiotics, stressors, nutrient sources) was found to be better predicted by our model. The lack of predictive power for some of the conditions could be due to the perturbation not being strong enough, lack of conservation of conditionally essential genes and/or incorrect prediction of deleterious effects of variants. It is however unlikely that variant effect predictions are the major source of error since our performance for strains that are more phylogenetically distant from the K-12 reference is equally good. We even observed a marginal improvement in predictive power when restricting the analysis to the 100 most phylogenetically distant strains from the reference (Supplementary figure 4). Weighting the contribution of the conditionally essential genes to each condition also improved the predictive power, especially for well-predicted conditions (Supplementary figure 4).

**Figure 4.**
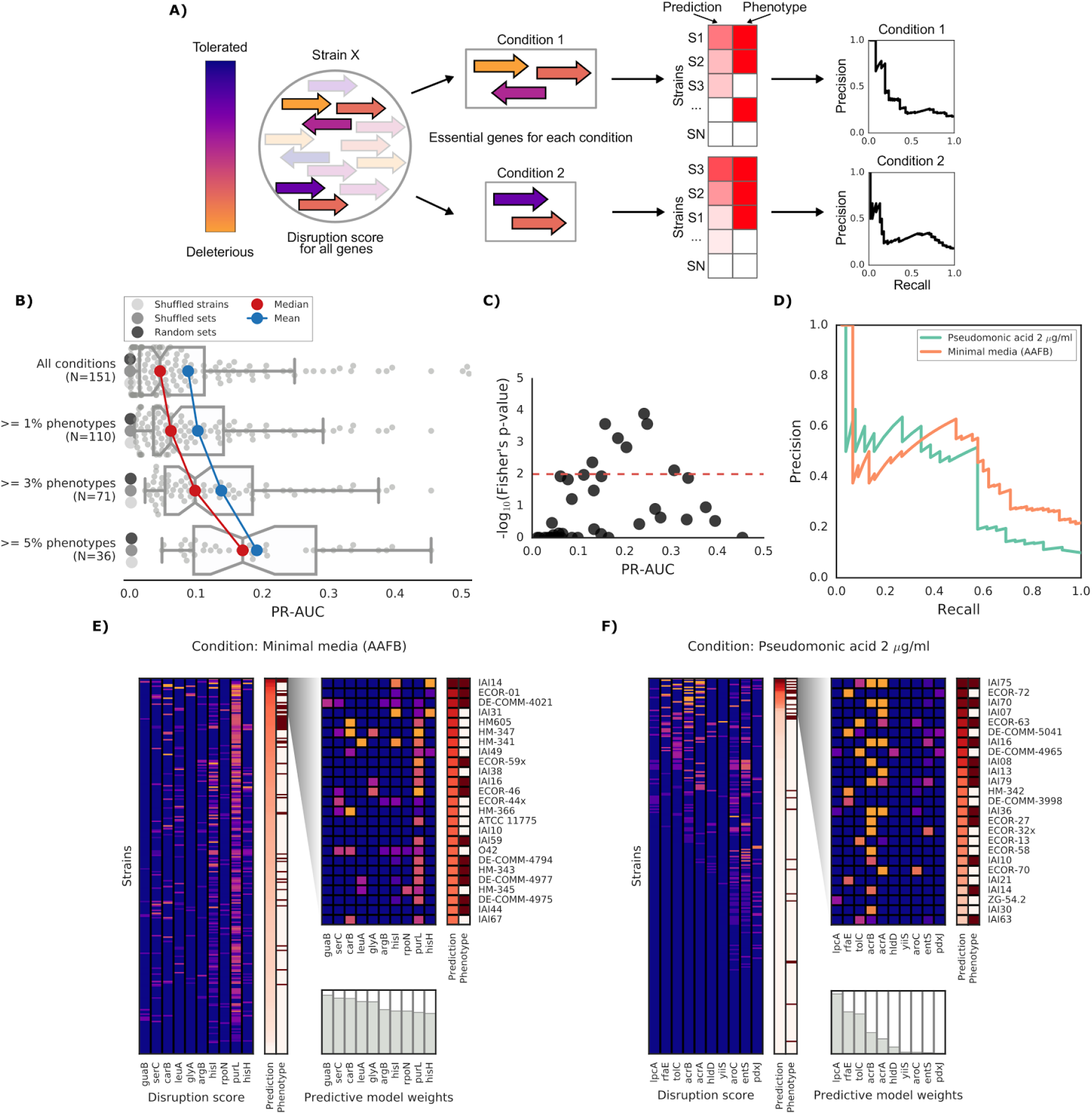
Prediction of growth-defect phenotypes in the *E. coli* strain collection. **(A)** Schematic representation of the computation of the prediction score and its evaluation; for each condition the predicted score is computed using the disruption score of the conditionally essential genes. The score is then evaluated to the actual phenotypes through a Precision-Recall curve. **(B)** Higher predictive power for conditions with higher proportion of growth phenotypes. For each condition set, the median and mean PR-AUC (area under the curve of the Precision-Recall curve) across conditions is compared to three randomisation schemes; one involving shuffling of strains (1’000 repetitions), one involving shuffling of conditionally essential genes (1’000 repetitions) and one using random genes as members of the conditionally essential gene sets (1’000 repetitions). The median PR-AUC and mean absolute deviation of the randomisations are reported across all conditions. **(C)** Genome-wide gene associations are in agreement with our predictive score; the enrichment (expressed as − *log*_10_ of a Fisher’s exact test P-value) of conditionally essential genes in the results of the gene association analysis is significantly higher in conditions with higher PR-AUC. **(D-E)** Detailed example on the computation and evaluation of the predicted score on two conditions. **(D)** Precision-Recall curve for the two example conditions. **(E-F)** For each condition, the gene disruption score for the conditionally essential gene across all strains is reported, together with the resulting predicted conditional score and actual binary phenotypes (pale red: healthy and red: growth defect). Strains are sorted according to the predicted conditional score, while genes are sorted according to their weight in the predictive model; only the top 10 conditionally essential genes are shown. The inset reports the disruption score, predicted score and actual phenotypes for the top 25 strains.

To independently validate our predictive models, we carried out a GWAS analysis based on genes presence/absence and the growth phenotypes. Consistent with the validity of our models, we found that for conditions we predict with higher confidence (PR-AUC >= 0.1), there is a significant overlap between the K-12 genes predicted to be essential in the condition and the genes associated with poor growth by the association analysis (Figure 4C, Fisher’s exact test, p-value 0.005).

We further examined two well predicted conditions (PR-AUC > 0.35), pseudomonic acid 2 μg/ml (the antibiotic mupirocin) and minimal media with the addition of aminoacid solutions, to inspect our predictive model (Figure 4E-F). Both conditions showed an enrichment of strains with growth defects at high predicted scores (GSEA p-values of 0.001 and < 10^−6^, respectively), which is a common property of conditions with higher PR-AUC (Supplementary figure 4). The reference K-12 strain harbors many conditionally essential genes in minimal media (181), providing an example for which growth phenotypes are well predicted from deleterious effects across a large number of genes in different strains. In contrast, pseudomonic acid is a condition with few (10) conditionally essential genes, thus making it easier to pinpoint single disrupted genes as causal for the phenotype. In these examples, the “healthy” strains that have a high predicted score, and are therefore incorrectly predicted as “sick” by the model, represent examples where conditional gene essentiality may not be conserved, or where compensatory mutations might be confounding the computation of the gene disruption score. For example, of the 25 strains with highest predicted score in pseudomonic acid, 13 have been misclassified by the model, 7 of which had strong disruption scores in either *gmhA* or *rfaE*, two genes that when deleted in K-12 cause a strong growth defect under pseudomonic acid. Only 1 of the correctly predicted strains showed a mutation in one of those two genes (IAI36), suggesting that these two genes might be conditionally essential only in K-12. Another example of incorrect predictions involves two strains (ECOR-27 and ECOR-58) that share very similar disruption score profiles for the conditionally essential genes in pseudomonic acid (Figure 4E), but only ECOR-27 exhibited a growth defect in this condition. Both strains harbor a single nonsynonymous mutation in *acrB* (E414G for ECOR-27 and I466T for ECOR-58), which, in both cases, is predicted as highly deleterious by the SIFT algorithm. Changes in conditional essentiality or epistatic effects are possible explanations for this misclassification, and therefore mapping and incorporating this information in our models can significantly benefit predictions in the future.

### Experimental validation of predicted causal variants

Mechanistic models of the impact of genetic variants on the phenotype can be used to directly predict the causal variants and implement genetic therapies to correct growth phenotypes. We tested this by ranking the mutated conditionally essential genes in each condition according to their predicted ability to rescue growth phenotypes (Figure 5A and Methods). Several genes are predicted to restore growth phenotypes across many strains and conditions (Figure 5B). Among them are the components of the AcrAB-TolC multidrug efflux pump, which were predicted to restore growth in up to ~1000 condition-strain pairs, would their disruption be reverted (1012 *acrB*, 494 *acrA* and 517 *tolC*, respectively); this finding reflects the importance of this efflux system in drug resistance (Li, Plésiat, and Nikaido 2015). We selected 8 genes to experimentally verify our predictions, including genes involved in either drug resistance or auxotrophic growth: two members of the AcrA-AcrB-TolC multidrug efflux pump (*acrA* and *acrB*), the peptidoglycan-degrading enzyme Slt, the first two enzymes in the L-proline biosynthetic pathway (ProA and ProB), the uridylyl transferase GlnD and the regulatory genes of the superoxide response (soxS and *soxR*). We then selected conditions for which the introduction of the reference allele in the target strains is predicted to restore growth, and controls where no improvement in phenotype is predicted (Methods). We also introduced the plasmid expressing the reference allele in the reference strain as a negative control and in the deletion strain from the KEIO collection as a positive control. Overall, we observed a high correlation between replicates of this experiment (Pearson’s r: 0.94, Supplementary figure 5). In total, we tested 64 strain-condition combinations, detecting a significant difference (p-value 8.4E-11, two-sided t-test) in colony size between strain-condition pairs in which we had predicted growth will be restored (N=14) versus ones we had not (N=50) (Figure 5C and Methods). This validated the effectiveness of our predictive models and more generally confirmed that genetic variants can be used to predict causal mutations and prioritise strategies for reverting/modifying phenotypes.

**Figure 5.**
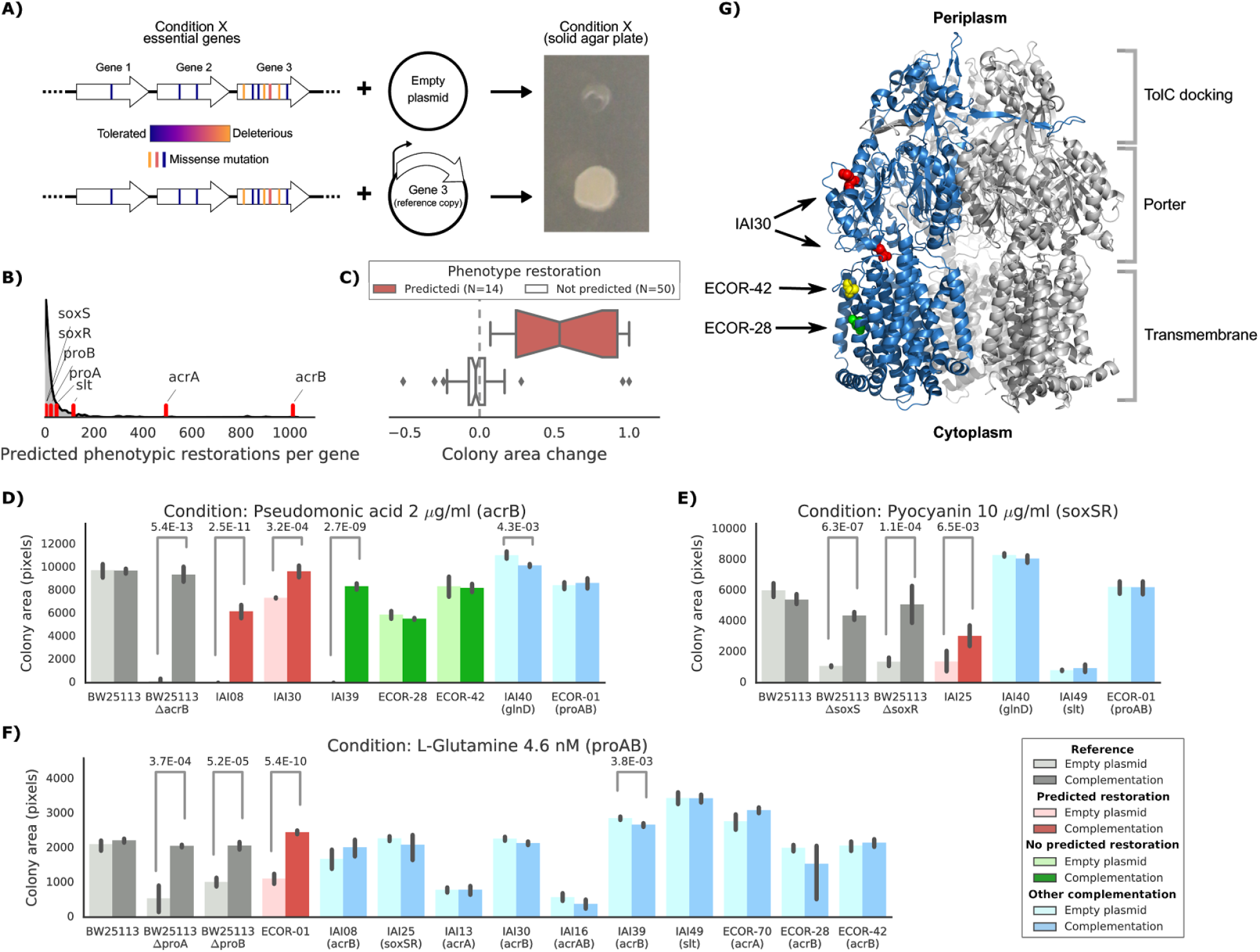
Experimental confirmation of the predicted phenotype-causing genotypes. **(A)** Schematic representation of the experimental approach. For each target condition and strain with a growth defect, we introduced a plasmid expressing the reference copy of the gene whose deleterious variants are predicted to give rise to the observed phenotype. **(B)** Distribution of the number of predicted restored phenotypes per gene; red stripes indicate the genes that were experimentally tested. (C) Growth change between the target strain with the empty plasmid against the one expressing the reference copy of the target gene (complementation). Strains for which a change in phenotype is predicted are compared to those where no change is predicted. **(D-F)** Detailed representation of the results of the complementation experiment in three conditions. Mean colony area and the 95% confidence intervals are reported. Significant differences (t-test p-value < 0.01) between colony area of the strains with the empty and complemented plasmids are reported. “Other complementation” reports strains expressing a different gene than the focal one, indicated between parenthesis. **(G)** Cartoon representation of AcrE3 3D structure (PDB entry: 2dr6). Only one of the three monomers is highlighted in blue; colored spheres represent known non-synonymous variants in one of the strains complemented experimentally. Variants present in strains ECOR-42 and ECOR-28 are in the transmembrane domain, while both variants present in IAI30 are in the porter domain.

We investigated three conditions in more detail: pseudomonic acid 2 μg/ml with an *acrB* complementation, pyocyanin 10 μg/ml (a toxic secondary metabolite produced by *Pseudomonas aeruginosa)* with a *soxSR* complementation and the L-glutamine aminoacid as nitrogen source with a *proAB* complementation (Figure 5D-F & Supplementary figure 5 for all conditions and strains). In all cases, strains harboring the gene(s) predicted to restore the phenotypes grew significantly better than strains carrying the empty plasmid (t-test p-value < 0.01). In contrast, none of the strains harboring a different complementation gene showed a significant increase in colony size. In fact, in two cases a significant, although slight decrease was observed, presumably due to toxic effects of overexpressing the target gene (strain IAI40 expressing *glnD* in pseudomonic acid and strain IAI39 expressing *acrB* in L-Glutamine). In one case we detected an unexpected increase in colony size for one of the strains where we hadn’t predicted any change (strain IAI39 expressing *acrB*, Figure 5D). This strain encodes a single nonsynonymous variant in *acrA* (T104A), which is predicted to be neutral (SIFT p-value > 0.05 and FoldX ΔΔG < 0.46 kcal/mol). We hypothesize that the original growth phenotype is either due to this incorrectly classified variant or due to another variant that indirectly acts on the expression of this efflux pump.

Of the 5 strains expressing the reference allele of *acrB* tested in pseudomonic acid, three harbored nonsynonymous variants in their chromosomal copy. One of them (IAI30) was predicted to increase its colony size after the complementation, whereas the other two (ECOR-42 and ECOR-28) were not predicted to be affected by the complementation (Figure 5D). We mapped those nonsynonymous variants to the three-dimensional structure of AcrB (PDB entry: 2dr6, Murakami et al. 2006) to inspect their potential impact on the protein function (Figure 5G). Both ECOR-42 and ECOR-28 harbor a single nonsynonymous variant (A915D and T1013I, respectively) in the transmembrane domain of the protein; both variants are predicted to be deleterious (SIFT, p-value ~0.01). Strain IAI30 on the other hand carries two nonsynonymous variants: E567V and H596N, both located in the AcrB “porter” domain, and more specifically in the PC1 subdomain, which is involved in the entry of the ligand that will be then extruded by the efflux pump (Murakami et al. 2006; Seeger et al. 2006). Since the E567V variant is predicted to be even more deleterious (SIFT, p-value ~0.001) than the variants present in the other two strains, we presume that this variant impairs the fundamental drug uptake function of the AcrAB-TolC multidrug efflux pump, while the other variants might be less deleterious for the pump’s function. This example shows how mechanistic interpretations of the impact of genetic variants can direct insights into the emergence of fitness defects and suggest potential gene therapeutic strategies, down to the level of the single genetic variant.

## Discussion

We have assembled an *E. coli* strain collection for genotype-to-phenotype studies, which, in its current state, comprises joint genetic and phenotypic information for 696 strains. We observe only weak to no correlation between the genotype and phenotype distances. This together with the large diversity of phenotypes detected even for strains deriving from evolutionary experiments, signifies that small differences in the genome can create large phenotypic variance.

To quantitatively assess the function of each reference gene in each strain, we computed a gene disruption score based on predictions of the impact of nonsynonymous substitutions to the protein structure/conservation and to gene loss. For this purpose, we used the pre-computed predicted consequences of all possible genome variants, using structural and evolutionary information. Those predictions have been made available online (http://mutfunc.com, Wagih et al., unpublished) where others can query any *E. coli* genome variants.

There are limitations to the disruption score used here. We evaluated the impact of individual and independent variants relative to a single reference (the K-12 genome) and we therefore didn’t take into account epistatic mutations within and between each gene. Nevertheless, the gene disruption score shows expected properties such as lower disruption of essential genes. In addition, genes within the same functional group (e.g. protein complex) tend to be co-disrupted across strains. Similar patterns of co-gain/loss of genes belonging to a functional unit have been observed when comparing genomes of different species (Pellegrini et al. 1999). This is thought to be due to changes in selection pressure whereby the reduced selection for the function carried out by a complex would result in the eventual loss of all of the genes of the complex. Our results suggest that the same process occurs at shorter evolutionary timescales and can therefore be detected even at the species level. Overall, this finding opens up the door for combining sequencing of large bacterial strain cohorts with function prediction models as the one presented here in order to map functional complexes.

We combined the gene disruption scores with the currently available reverse genetic screens in the reference individual to predict conditional phenotypes across 148 conditions. The impact of mutations on conditionally essential genes has an overall weak but significant predictive power, with 20-38% of conditions being well predicted by this approach. While it is possible that a fraction of the conditions is poorly predicted due to technical limitations (e.g. chemical concentration screened) or errors/limitations of the gene disruption scores, these results strongly suggest that the relative importance of each gene, and therefore gene function itself, may quickly diverge across strains. Previous work has shown that gene-specific phenotypes can diverge for around 17% of tested gene knock-outs across two very closely related *S. cerevisiae* laboratory strains (S288C and Σ1278b, Ryan et al. 2012). More strikingly, only 52% of essential genes were found to be conserved across 5 different human cancer cell lines (Hart et al. 2015). However, the genetic and cell state differences across the *E. coli* strains cannot easily be related to the differences between cancer cell lines or closely related S. *cerevisae* strains. Additional studies are required to conclusively demonstrate that gene function and conditional essentiality diverges rapidly across *E. coli* strains.

We further demonstrated how the predictive model used here can be used to identify the causal variants and suggest genetic strategies for restoring growth defects. These models highlighted genes that are very often predicted to restore growth across many conditions and strains. In particular, the AcrAB-TolC pump emerged as a complex that is often perturbed across evolution and as predicted, restored growth when complemented.

There are several ways by which the predictive approach applied here could be further improved. For example, the model can be further expanded to include the impact of non-coding variants or accessory genes not present in K-12. Reliable predictors for assessing the impact of non-coding variants on gene expression and translation will need to be developed and included in the model. A better understanding of the function of the accessory genes would also be needed for predicting when gene loss events are complemented by the acquisition of new genes, either through recombination or horizontal gene transfer (HGT). Finally, additional work will be needed to take into account gain-of-function mutations or epistatic interactions, whereby the effect of variants depends on the genetic background of the individual.

Following the example of previous genetic reference panels (Ayroles et al. 2009; Liti et al. 2009; Bennett et al. 2010; Atwell et al. 2010; 1001 Genomes Consortium 2016; Cancer Genome Atlas Research Network et al. 2013; 1000 Genomes Project Consortium et al. 2015), we believe that the present strain collection and associated data will serve as a growing resource to researchers interested in studying other aspects of basic and bacterial biology. Any measurements or models added to these strains will amplify the benefit for the entire research community, moving us closer to the ultimate goal of understanding of how genetic variation translates to differences among individuals.

## Acknowledgements

We are particularly grateful to the various people providing us with many of the strains of the *E. coli* genetic reference panel, specifically (in alphabetical order): Alfredo G. Torres, Catharina Svanborg, David Clarke, Erick Denamur, Ewa Bok and Pawel Pusz, Isabel Gordo and Lilia Perfeito, Jorg Weinreich and Peter Schierack, KC Huang, Lisa Nolan, Mark Goulian, Mathew Upton, Olin Silander, Richard Lenski, Scott Hultgren and Wanderley Dias da Silveira. We thank Amanda Miguel for helping in the phenotypic screen. We also thank the EMBL Gene Core, and especially Rajna Hercog and Vladimir Benes for the support in genome sequencing. We thank Ewan Birney, Oliver Stegle, KC Huang and Zam Iqbal for critical reading of the manuscript.

This work was partially supported by the Sofja Kovaleskaja Award of the Alexander von Humboldt Foundation to ATy and a grant from the Fondation pour la Recherche Médicale (Equipe FRM 2016, DEQ20161136698) to ED.

All the authors declare no conflict of interest.

## Contributions

PB and ATy conceived the study, ATy, LHD and MG designed the phenotypic screening, LHD, ATe and MG carried out the phenotypic screening, ATe sequenced part of the strains, MG and JACV analysed the sequencing data, OW and MG precomputed the predicted impact of nonsynonymous substitutions, MG designed the strain collection, analysed the phenotypic data and applied the predictive models, MG, AK and ATy designed the follow-up experiments, AK carried out the follow-up experiments, AK and MW assembled the strain collection, OC and ED provided part of the strain collection, OC, ED and MG verified the consistency of the genome sequences, MG, PB and ATy wrote the manuscript.

## Supplementary figures

**Supplementary Figure 1. Related to Figure 1.**
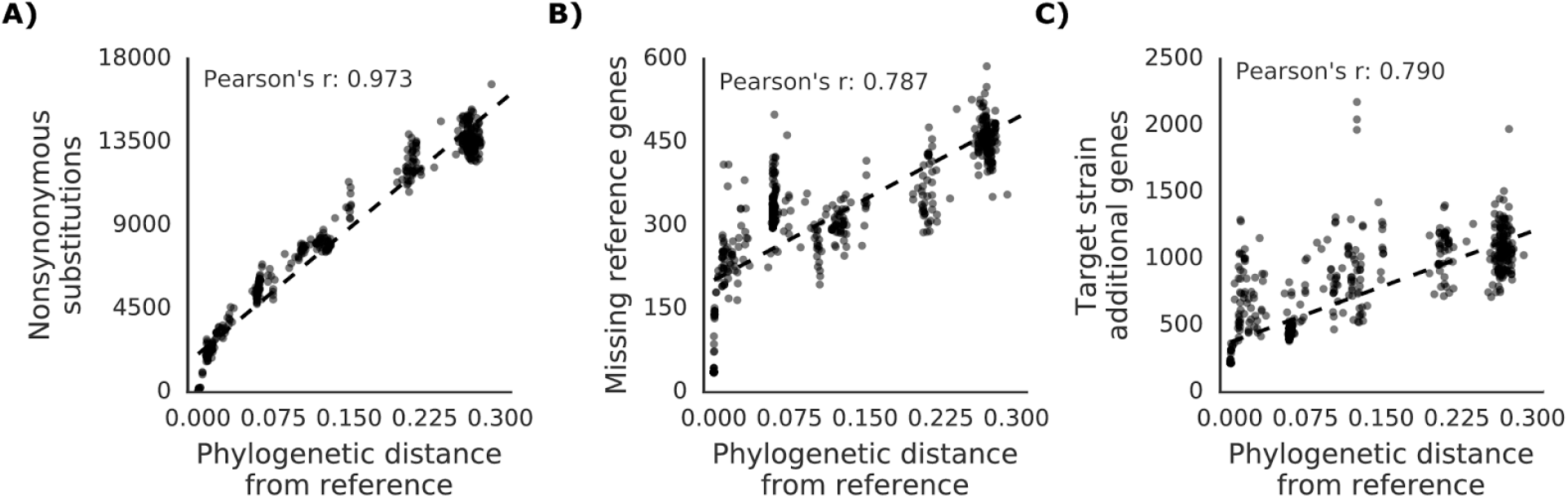
Correlation between phylogenetic distance and genetic variants of the members of the strain collection. Phylogenetic distance is derived from the core genome alignment of all the strains. **(A)** correlation with the number of nonsynonymous substitutions. **(B)** correlation with the number missing reference genes. (C) Correlation with the number of genes present in the target strain and not in the reference.

**Supplementary Figure 2. Related to Figure 2.**
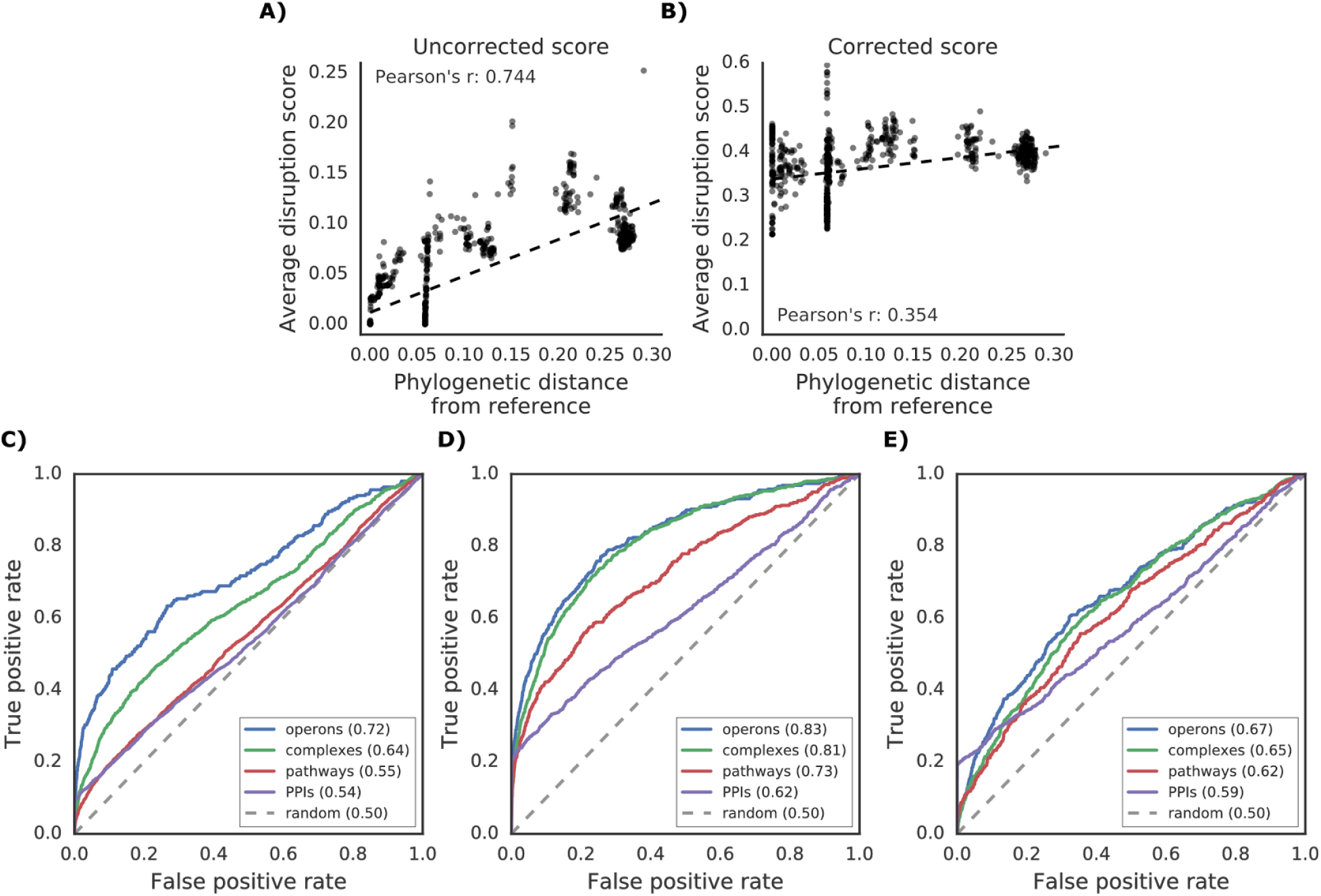
Additional properties of the disruption score. **(A)** Correlation between phylogenetic distance from the reference strain and average disruption across all reference genes. **(B)** Significant reduction in correlation after correction of the disruption score (see Methods). **(C-E)** Prediction of gene functional associations using disruption score profiles. **(C)** ROC curve using disruption score profiles across all genes. **(D-E)** Higher predictive power of the gene disruption profile when restricted to accessory genes, including **(D)** or not **(E)** the information about gene presence absence.

**Supplementary Figure 3. Related to Figure 3.**
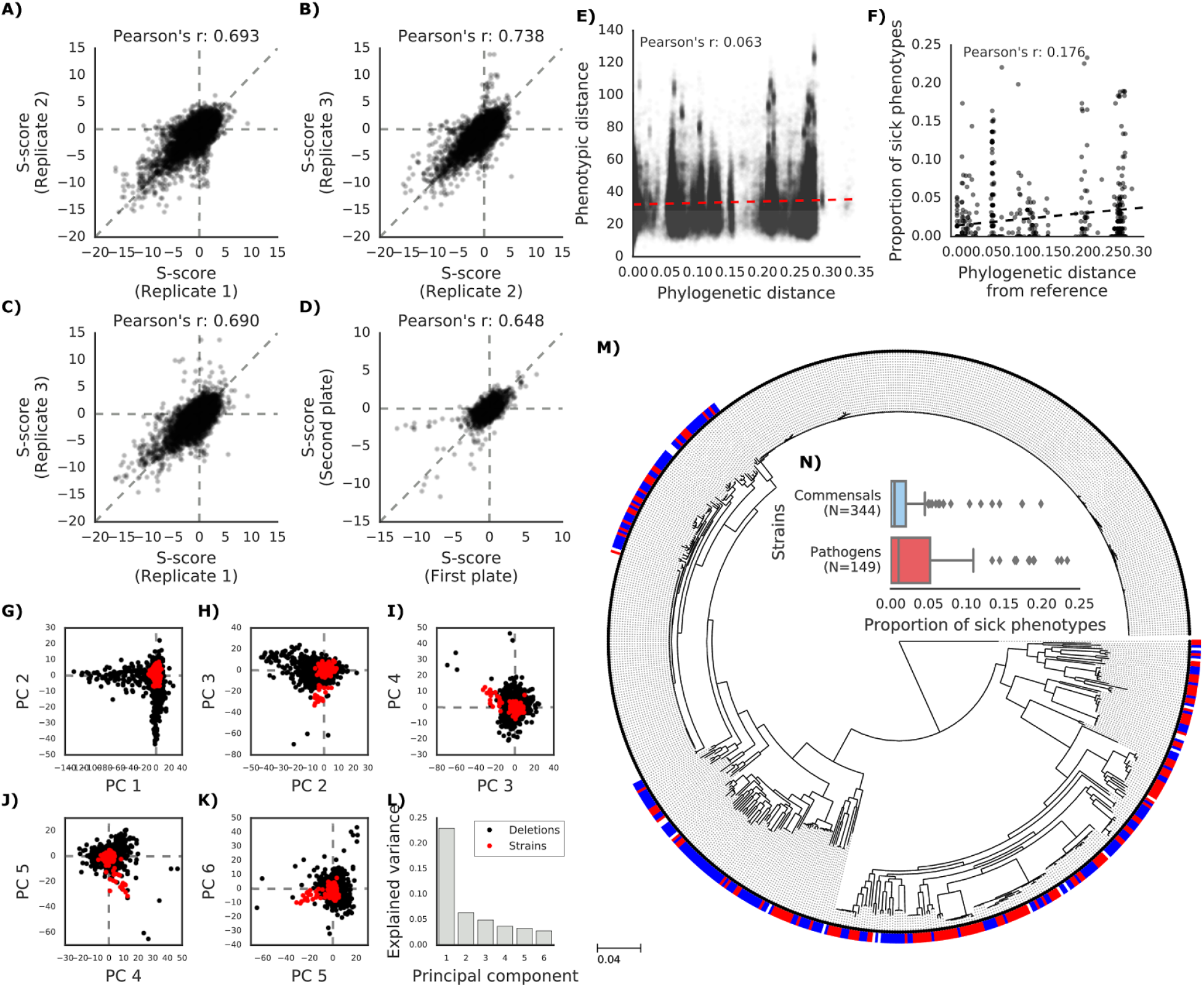
**(A-D)** Phenotypic measurements reproducibility. **(A-C)** S-scores for each replicate are compared against each other **(D)** S-scores comparison between strains present in multiple 1536 plates. **(E)** Linear correlation between phylogenetic and phenotypic distance between each pairwise strain combination. **(F)** Linear correlation between phylogenetic distance from the reference strain and proportion of sick phenotypes over tested conditions. **(G-L)** PCA (Principal Component Analysis) of the joint phenotypic measures for the chemical genomics and the *E. coli* strain collection. Only the first 6 principal components and their relationships are reported. (I) Proportion of total explained variance for the first 6 dimensions. **(M-N)** Differences in the proportion of sick phenotypes between commensal and pathogenic strains. **(M)** Core genome SNP tree of the members of the strains collection; blue and red boxes indicate commensal and pathogenic strains, respectively. **(N)** Proportion of sick phenotypes over the tested conditions for each strain divided according to their broad phenotype (commensal or pathogen), showing a moderately significant difference (Cohen’s d: 0.658).

**Supplementary Figure 4. Related to Figure 4.**
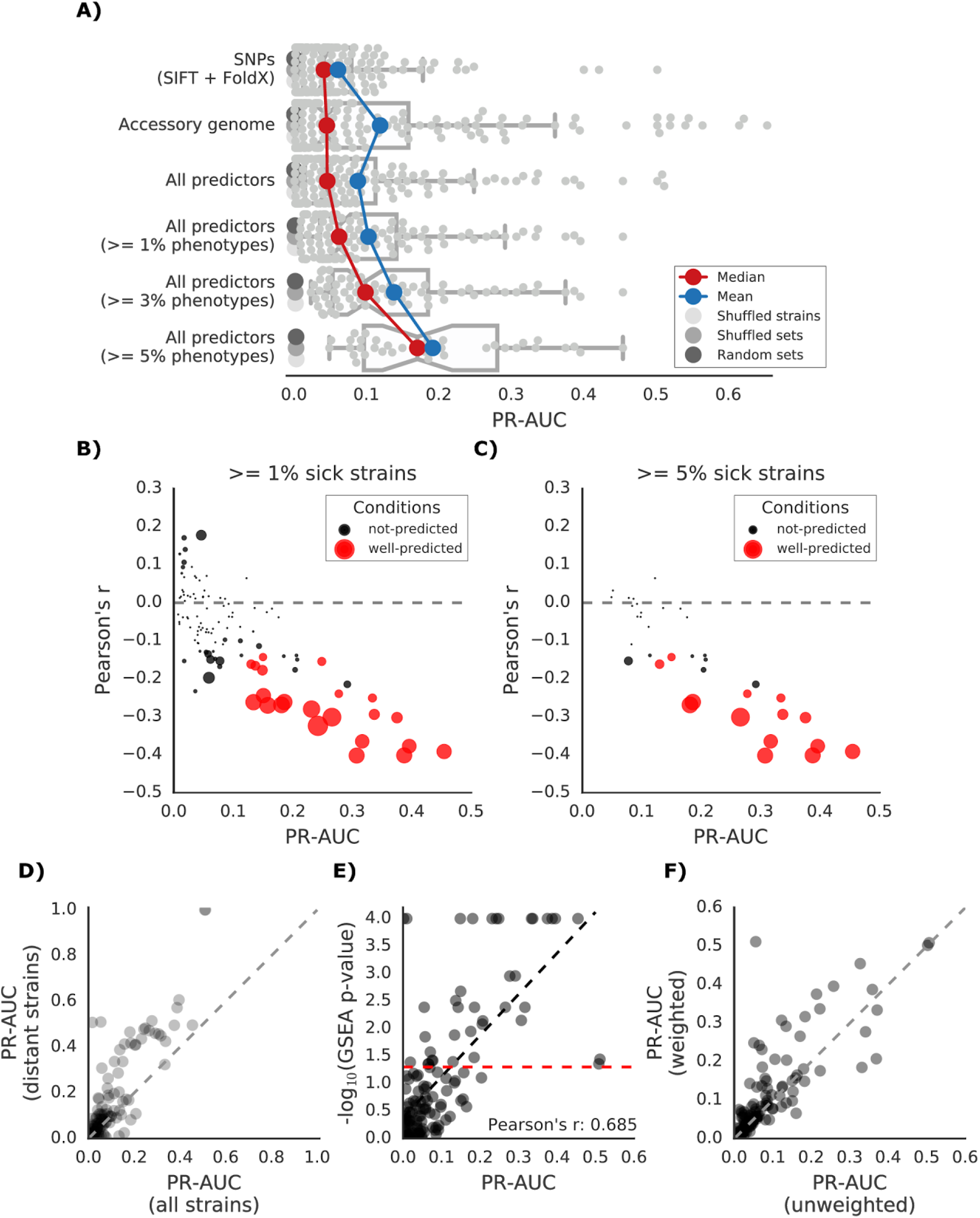
**(A)** Influence of the different predictors of the impact of mutations on each condition PR-AUC. “SNPs” indicates single nucleotide variants only, “Accessory genome” gene presence-absence patterns, “All predictors” the combination of both. **(B-C)** Proportion of well-predicted conditions (PR-AUC >= **0.1** and Pearson’s FDR-corrected p-value <= 0.01) over total conditions with at least 1% and 5% sick strains. Marker’s size is proportional to the −*log*_10_ of the FDR-corrected p-value. **(D)** Improvement in prediction performance when restricting the analysis to the 100 strains most distant from the reference (K-12). **(E)** Conditions with higher predictive power (measured as PR-AUC) also have an enrichment of sick strains at the top of the predicted score, as measured by the Gene Set Enrichment Analysis (GSEA); sick strains are used as “gene sets”. A pseudocount of 10^−4^ has been added to the GSEA p-values. (F) Prediction performance improves when using the weighting scheme to account for conservation of gene essentiality, especially for well-predicted conditions.

**Supplementary Figure 5. Related to Figure 5.**
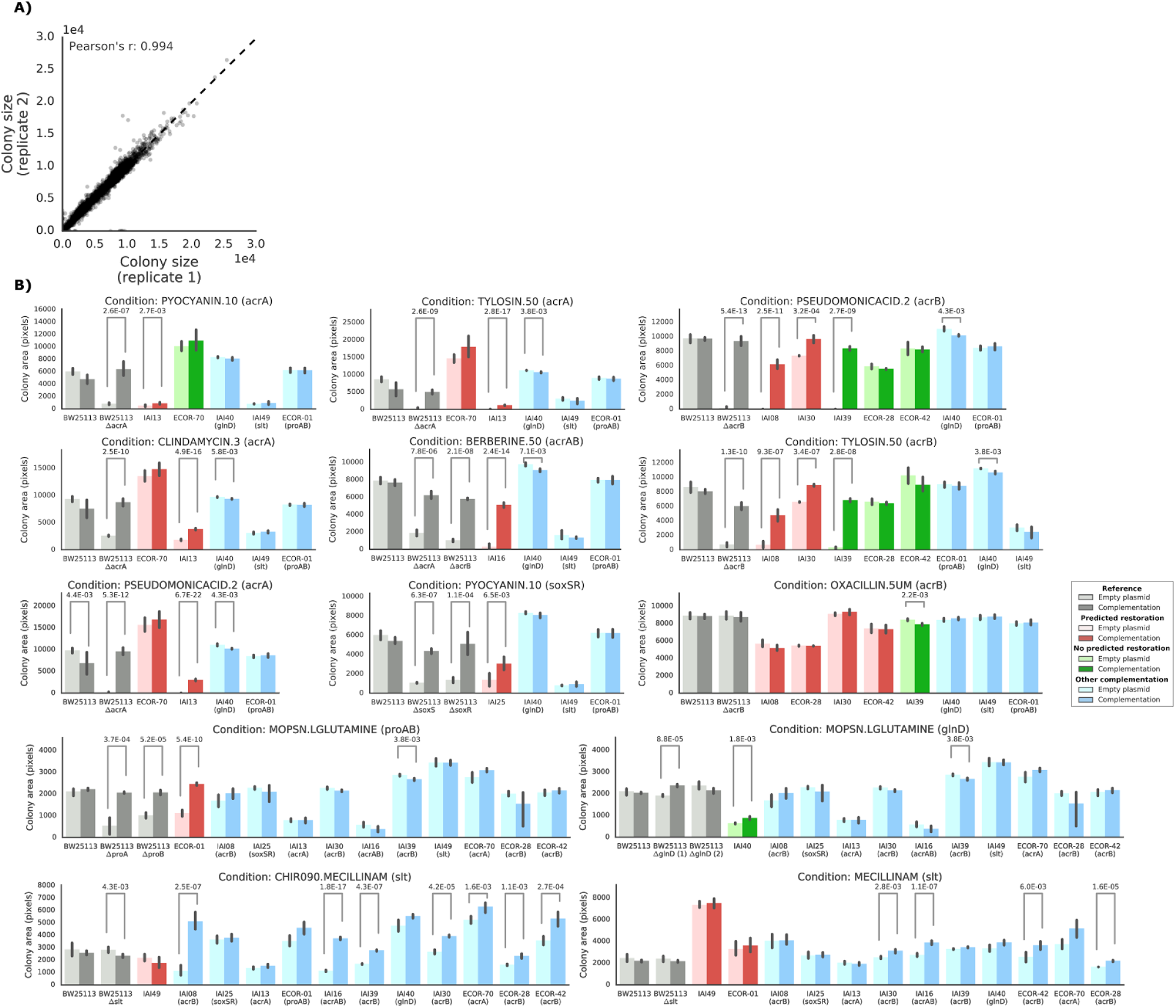
**(A)** Replicability in colony size (measured in pixels) after outer frame and spatial correction between the two replicates. **(B)** Overview of the results of the complementation experiments. Mean colony area and the 95% confidence intervals are reported. Significant differences (t-test p-value < 0.01) between colony area of the strains with the empty and complemented plasmids are reported. Conditions CHIR090.MECILLINAM, MECILLINNAM and OXACILLIN.5UM have been excluded from further analysis, as the deletion strain from the KEIO collection (positive control) does not show a sick phenotype with an empty plasmid.

## Materials and methods

### Genome sequencing, assembly and annotation

Strains whose genome sequence was not yet available were sequenced using various Illumina paired-end platforms (Supplementary table 1). The resulting sequencing reads were quality checked using FastQC version 0.11.3, and optionally contaminating sequencing adapters were removed using seq_crumbs version 0.1.9. Reads were assembled with Spades (Bankevich et al. 2012) version 3.5.0, using different k-mer sizes according to reads length and with the “careful” option to reduce assembly errors; contigs below 200 base pairs were excluded. Resulting assembled contigs were annotated for coding genes, ribosomal RNAs and tRNAs using Prokka (Seemann 2014) version 1.11. Strains not belonging to the *E. coli* species were excluded from subsequent analysis after being highlighted by Kraken (Wood and Salzberg 2014) version 0.10.5. When available, typing information was used to spot incorrect genome sequences due to culture contaminations or other factors; known strains typing was compared to the ones predicted from the genome sequence using mlst version 2.8. Strain names were amended when possible (see the “Notes” column in Supplementary table 1). The ECOR collection was carefully checked for inconsistencies, as it is well known that different “versions” of this collection are circulating in the scientific community (Johnson et al. 2001; Clermont, Gordon, and Denamur 2015). The genome sequences were further checked for duplicated genomes: strains with highly similar genomes but highly divergent phenotypes (phenotypes S-score correlation below 0.6) were removed. Highly similar genomes were defined as those genomes whose distance was found to be below 0.001, as measured by mash, version 1.1 (Ondov et al. 2016).

### SNP calling and annotation

Due to the variability in sequencing technologies or lack of the original reads for the already sequenced strains in the collection (373), SNPs were called through a whole genome alignment between each strain and the genome of the reference individual (*Escherichia coli* str. K-12 substr. MG1655, RefSeq accession: NC_000913.3, strain collection identifier NT12001), using ParSNP (Treangen et al. 2014) version 1.2. Repeated regions in the reference genome were highlighted and masked through nucmer (Kurtz et al. 2004) version 3.1 and Bedtools (Quinlan and Hall 2010) version 2.26.0. SNPs were then phased, and annotated using SnpEff (Cingolani et al. 2012) version 4.1 g.

### Pangenome analysis

Genes present in the reference individual but absent in each strain were highlighted by computing hierarchical orthologous group using OMA (Altenhoff et al. 2013) version 1.0.6. Each strain was re-annotated using Prokka (Seemann 2014) version 1.11 to harmonize gene calls.

### Phylogenetics

Strains phylogenetic tree was computed using a single ParSNP (Treangen et al. 2014) analysis, which uses the regions of the reference genome that are sufficiently aligned across all strains as an input for FastTree (Price, Dehal, and Arkin 2010) version 2.1.7. The tree was visualized using the ete3 library (Huerta-Cepas, Serra, and Bork 2016) version 3.0.0.

### Computation of all possible mutations and their effect

The impact of all possible nonsynonymous substitutions on the reference individual has been precomputed to speed up the lookup process. Functional impact of nonsynonymous substitutions has been computed using SIFT (Ng and Henikoff 2001) version 5.1.1. Structural impact of nonsynonymous substitutions has been computed using FoldX (Guerois, Nielsen, and Serrano 2002) version 4; both 3-D structures present in the PDB database and homology models were used. Homology models were created using ModPipe (Pieper et al. 2014) version 2.2.0. Water accessibility of all 3-D structures was computed using FreeSASA (Mitternacht 2016) version 1.1. Conversion from PDB to Uniprot residues coordinates was derived from the SIFTS (Velankar et al. 2013) database. All the precomputed impacts of all possible nonsynonymous substitutions are available through the mutfunc database (http://mutfunc.com, Wagih et., al, unpublished).

### Computation of the disruption score

For each strain and each protein coding genes we have computed the overall impact of all nonsynonymous and nonsense substitutions, in a similar approach as the one used for *Saccharomyces cerevisiae* (Jelier et al. 2011). The output of each predictor (a deleterious probability for SIFT and a ΔΔG value for FoldX) has been converted to the probability of the substitution being neutral *P*(*neutral*). Mutations with known impact on the reference individual have been downloaded from Uniprot (UniProt Consortium 2015) (N=3673) and used to derive such conversion; since only 580 mutations are reported to have a neutral impact, we added all observed nonsynonymous variants affecting known essential genes (as reported in the OGEE database, Chen et al. 2016) to the list of tolerated mutations. The distribution of the negative natural logarithm of the SIFT probability (plus a pseudocount equivalent to the lowest observed SIFT probability) for all the 6460 mutations was binned and the *P*(*neutral*) was computed as the proportion of tolerated mutations over the total number of mutations in each bin. A logistic regression curve was then fitted to the binned distribution to derive the conversion between the SIFT probability and *P(neutraĩ)*. For FoldX we used a similar approach, but using the computed ΔΔG value. The fitted logistic regression curves resulted in the following *P*(*neutral*) functions:

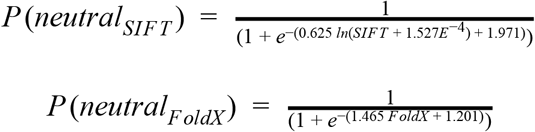

The *P*(*neutral*) value attributed to nonsense mutations was assigned through heuristics: if the new stop codon was found within the last 16 residues of the protein it was given a *P*(*neutral*) value of 0.99, reflecting its unlikelihood of disrupting the function of the protein, 0.01 otherwise. Losing a start or a stop codon was given a *P*(*neutral*) value of 0.01, as they are very likely to impair protein function.

We gave a *P*(*neutral*) value of 0.01 to those genes that were found to be present in the reference individual but absent in the target strain, reflecting the fact that their function is most likely to be absent from the target strain.

We inferred the probability that each gene had its function affected by the ensemble of the substitutions in each strain by computing a disruption score, equivalent to the *P*(*AF*) (probability of the function being affected) used in S. *cerevisiae* study (Jelier et al. 2011).

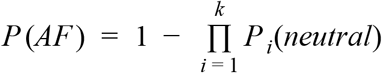

Where *k* is the ensemble of nonsynonymous and nonsense substitutions observed in each gene. When both FoldX and SIFT predictions were available for a given substitution, we used the SIFT prediction only. Variants with relatively high frequency (>= 10%) in the strains collection were not considered, as well as those reference genes that are absent in a significant number of strains (>= 10%), as we reasoned that they are very unlikely to be deleterious given their high observed frequency. Given that many strains of the collection are closely related (e.g. strains derived from the LTEE collection), we clustered them based on phylogenetic distance before applying the filtering. We also didn’t consider variants and absent reference genes shared by all members of the LTEE strain collection, as those variants are present in the collection founder strain and therefore unlikely to affect the evolved clones phenotypes. Disruption scores for all proteins across all strains can be found in Supplementary material 2.

### Use of the disruption score as a functional association predictor

We used the proteins pairwise Pearson’s correlation of disruption scores as a predictor of genes functional associations. We used four benchmarking sets: operons, as derived from the DOOR database (Mao et al. 2014), protein complexes and pathways derived from the EcoCyc database (Keseler et al. 2013), and protein-protein interactions derived from a recent yeast two-hybrid experiment (Rajagopala et al. 2014). The distribution of the disruption score correlation for each pair of related genes was compared against the same number of gene pairs randomly drawn from all reference genes. We also assessed the predictive power of the disruption score correlations by drawing a receiving operator characteristics curve (ROC) across each correlation threshold, using the scikit-learn library (Pedregosa et al. 2011) version 0.17.1.

### Strains phenotyping

The strains phenotypes were measured in a similar way as the *E. coli* reverse genetic screen (Nichols et al. 2011). The strain collection was plated in three solid agar plates, each one containing 1536 single colonies, so that each strain was plated at least four times in each plate, each time with different neighboring strains. For each condition we prepared three replicates (each one using a different source plate to reduce batch effects) with the concentrations indicated in Supplementary table 2 and the addition of the so-called Congo-red solution, which contains the Cosmos dye (CAS number 573-58-0) and Coomasie brilliant blue R-250 (CAS number 6104-59-2). The solution stains colonies when biofilm is being developed. Plates were stored in darkness at room temperature, unless otherwise required by the specific condition (i.e. higher temperature), and photographs of the plates were taken until colonies were found to be overgrowing into each other. Most of the conditions (197) were tested at the same time and under the same laboratory conditions as the KEIO KO collection (Herrera-Dominguez et al., unpublished).

A series of colony parameters were extracted from each photograph, using Iris (Kritikos et al. 2017) version 0.9.4.71: colony size, opacity, roundness and color intensity. The most appropriate time point for each condition was determined by imposing a restriction on median colony size; between 1900 and 3600 pixels for the first two plates, and between 1300 and 3600 for the last plate, which contained the strains derived from evolution experiments, which tend to grow less than the natural isolates. The time points passing the first threshold were then sorted by the proportion of colonies with high roundness (> 0.8), which is indicative of the overall quality of the plate, proportion of colonies over the minimum median colony size threshold, the spread of the colony size distribution (the lower the better), and mean colony size correlation across replicates.

A series of additional quality control measures were taken on the colony parameters. In order to remove systematic pinning defects, colonies appearing to be missing (colony size of zero pixels) in more than 66% of the tested plates were removed, unless all the internal replicates were found to be missing. Colonies with abnormal circularity were removed, as they were mostly due to incorrect colony recognition by the software: colonies with size below 1000 pixels and circularity below 0.5 and colonies with size above 1000 pixels and circularity below 0.3 were removed. Putative contaminations were spotted and removed through a variance jackknife approach: first, the size of two outermost rows and columns colonies was corrected to match the median of the rest of the plate, then for each strain, each of the four replicates inside the plate was tested whether it contributed to more than 90% of the total colony size variance. If so, the replicate was flagged as a contamination and removed. The same approach was repeated using colony circularity, with a variance threshold of 95%.

The final colony sizes were used as an input for the EMAP algorithm (Collins et al. 2006), with default parameters, in order to derive an S-score, which informs on the deviation of each strain from the expected growth in each condition. Final S-scores were quantile-normalized, and significant phenotypes were highlighted using a 5% FDR correction similar to the one used in the *E. coli* reverse genetic screen (Nichols et al. 2011), using the statsmodels library version 0.6.1. The phenotypic data is available in Supplementary material 1.

Phenotypic distance between strains was computed using the euclidean distance between the S-scores across all conditions, using the nadist library version 0.1.0. PCA (Principal Component Analysis) of the joint data from the K-12 chemical genomics and the *E. coli* strain collection was carried out by merging the two datasets over the shared conditions (conditions tested in both sets), using the scikit-learn library (Pedregosa et al. 2011), version 0.17.1. Since the PCA method doesn’t allow missing values, strains that were not tested in more than 60 conditions (over 161) were removed. The remaining missing values were imputed using the mean S-score across each condition.

### Computation of the conditional score and its assessment

Conditionally essential genes were derived for each condition of the *E. coli* reverse genetic screen overlapping with the conditions tested on the natural isolates collection. Mutants with a significant growth phenotype were considered to derive the list of conditionally essential genes. The conditional score for each strain, indicating the growth prediction, was computed as follows:

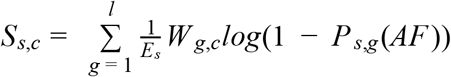

Where *s* and *c* represent the strain and condition, respectively, *g* each conditionally essential gene for condition *c* (with size *l*), and *E_s_* representing a correction term for the disruption score, in order to remove the effect of phylogenetic distance (Supplementary figure 2).

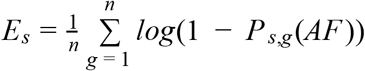

Where *n* represents all the reference genes. The term *W_g,c_* is used to weight the contribution of each conditionally essential gene to the conditional score, and it is computed as follows:

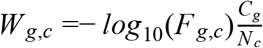

Where *F_g,c_* is the FDR-corrected p-value of gene *g* in condition *c*, *C_g_* is the number of conditions in the chemical genomics screen where gene *g* shows a significant phenotype, and *N_c_* the total number of tested conditions in the chemical genomics screen.

The conditional score was assessed by computing a Precision-Recall curve, whose area (PR-AUC) was used as a direct measure of the predictive power of the method; the growth phenotypes were considered true positives. The PR curve and AUC were computed using the scikit-learn library (Pedregosa et al. 2011) version 0.17.1. Three randomization approaches were used to generate control conditional scores: one using strains shuffling, one using conditionally essential gene sets shuffling, and one using random conditionally essential gene sets. Each randomization strategy has been used to generate 10,000 randomized scores, which were scaled to the actual one. The conditionally essential gene sets, the conditional score and the PR-AUC values are available in Supplementary material 2.

### Association of accessory genes with phenotypes

Accessory genes from the strains collection pangenome were computed from the harmonized genome annotations made by Prokka (Seemann 2014), using Roary (Page et al. 2015) version 3.6.1. The accessory genes were associated to each condition’s phenotypes using Scoary (Brynildsrud et al. 2016) version 1.4.0, with default parameters. Genes with corrected p-value (Benjamini-Hochberg) of association below 0.05 were considered significant. The enrichment of conditionally essential genes among the significant reference gene hits was assessed through a Fisher’s exact test, as implemented in the SciPy library, version 0.17.0.

### Systematic *in-silico* complementation of conditionally essential genes

The potential to restore growth phenotypes through the introduction of reference alleles was predicted systematically in each strain by changing the disruption score to zero in each conditionally essential gene, and reporting the change in the conditional score Δ*S_,c_* with respect to the maximum possible conditional score:

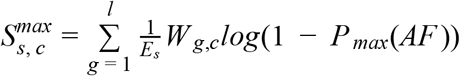

Where *P_max_*(*AF*) is the maximum disruption score observed across all genes and strains. Any Δ*S_s,c_* higher than 1% of 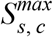 was considered as potentially able to restore a growth phenotype.

### Experimental complementation of predicted phenotype-causing genes

In order to experimentally verify our predictions, we introduced the reference (BW25113) gene in a low copy plasmid. For the *slt* gene, we used the available plasmid from the TransBac library (H. Dose and H. Mori, unpublished resource, Otsuka et al. 2015). For *acrA, acrB* and *glnD*, we used the available plasmid from the mobile plasmid library (Saka et al. 2005). We amplified *acrAB*, *soxSR*, *proAB* from BW25113 and ligated into pNTR-SD (the backbone plasmid for the mobile plasmid library). Deletions of *acrAB*, *soxSR*, *proAB* in the reference strain were made using the lambda red recombination approach (Datsenko and Wanner 2000). The resulting 7 plasmids and the 2 empty plasmid controls were introduced into BW25113 (negative control), in the deletion strains from the Keio collection or constructed by us (positive control) and in the targets strains. All resulting strains were pinned using a Singer Rotor robot in 10 different conditions, on 2 solid agar plates, so that each strain is pinned at least four times per plate. The plates were incubated at room temperature and multiple photographs were taken until colonies were found to be overgrowing into each other. Iris (Kritikos et al. 2017) version 0.9.7 was used to extract colony size from the pictures.

### Code, data and strains collection availability

The source code used to perform the analysis reported here and generate the figures is available as Supplementary material 3 and at the following URLs: https://github.com/mgalardini/screenings, https://github.com/mgalardini/pangenome_variation, https://github.com/mgalardini/ecopredict. Code is mostly based on the Python programming language, and using the following libraries: Numpy (Van Der Walt, Colbert, and Varoquaux 2011) version 1.10.4, SciPy version 0.17.0, Pandas (McKinney and Others 2010) version 0.18.0, Biopython (Cock et al. 2009) version 1.68, scikit-learn (Pedregosa et al. 2011) version 0.17.1, fastcluster (Müllner 2013) version 1.1.20, statsmodels version 0.6.1, PyVCF version 0.6.8, ete3 (Huerta-Cepas, Serra, and Bork 2016) version 3.0.0, Matplotlib (Hunter 2007) version 1.5.1, Seaborn (Waskom et al. 2016) version 0.7.1 and svgutils version 0.2.0.

Genomic and phenotypic data, as well as relevant information on how to obtain the members of the strain collection is available at the following URL: https://evocellnet.github.io/ecoref.

## Supplementary material

**Supplementary material 1:** s-scores, FDR corrected p-values and binary growth defect matrix of the E. coli phenotypic screening (869 strains across 214 conditions)

**Supplementary material 2:** disruption scores, conditional scores and prediction assessments.

**Supplementary material 3:** code used to analyze the genetic and phenotypic data, predict phenotypes and draw all the manuscript figures

